# Somatic variant detection in normal tissues from single-cell sequencing data

**DOI:** 10.64898/2026.06.10.731451

**Authors:** Rui Luo, Ziyi Wang, Jinzhuang Dou, Sravya V. Bhamidipati, Divya Kalra, Christopher M. Grochowski, HarshaVardhan Doddapaneni, Richard A. Gibbs, Ken Chen, Rui Chen, the Somatic Mosaicism across Human Tissues Network

## Abstract

A crucial advantage of single-cell sequencing (SCS) is its ability to identify somatic variants in individual cells, enabling phylogenetic analysis of cellular populations within bulk tissues. While identifying somatic variants in tumor tissues via SCS has become a common practice, doing so in normal tissues remains challenging due to the rarity of somatic variants in normal cells. To evaluate the feasibility of somatic variant calling from widely available single-nucleus RNA-seq (snRNA-seq) and single-nucleus ATAC-seq (snATAC-seq) data, we profiled a Cell-line mix of six HapMap samples prepared by the SMaHT consortium using 10x Genomics 5’ snRNA-seq (12k cells with 36k mean reads per cell) and snATAC-seq (11k cells with 14k median high-quality fragments per cell) for variant calling. PacBio long-read whole genome sequencing (WGS) data (109×) generated from individual cell lines were used as ground truth. Two computational tools, Monopogen and SComatic, were used for somatic variant calling from the SCS data. Monopogen achieved single nucleotide variant (SNV) detection accuracies of 93.30% in the snRNA-seq and 99.64% in the snATAC-seq data, both of which outperformed SComatic (74.35% and 94.29%, respectively). Monopogen also consistently detected somatic SNVs at cellular fractions as low as 0.5% (2.54% in snRNA and 0.81% in snATAC) in individual samples. Notably, snATAC-seq exhibited higher genomic coverage breadth and larger number of variants detected than snRNA-seq. While the SCS data have lower overall genome coverage than that of the bulk WGS, the single-cell level variant resolution allows Monopogen to assign variants to their cells of origin with over 80% accuracy in both RNA and ATAC modalities, thereby facilitating studies of clonal evolution and cell-type-specific mutagenesis. Other benchmarking methods were also evaluated (DeepVariant, Cellsnp-lite and Mutect2) for comparison. In conclusion, our study demonstrated the feasibility of performing reliable single-cell somatic mutation calling in a cell-line mixture and discussed the strengths and limitations of current computational methods when applied to normal tissues.

## Introductions

Somatic mutations accumulate throughout the human lifespan as a result of errors in DNA repair or replication, chromosome missegregation, or the activity of mobile elements^1–4^. Although most mutations are functionally neutral^5^, some can alter cellular phenotypes and contribute to disease. Insights into somatic variation have largely come from sequencing cancer genomes^6^, but more recent efforts, such as the Brain Somatic Mosaicism Network^7^, have revealed the presence of somatic mutations in normal tissues. Building on these initiatives, the NIH Common Fund established the *Somatic Mosaicism across Human Tissues (SMaHT) Network*^8^ to generate a reference catalogue of somatic variation from 150 donors across 19 non-diseased tissues, with a strong emphasis on advancing technologies for somatic mutation discovery.

Single-cell sequencing (SCS) offers a powerful means to interrogate somatic variation, in principle capturing all mutations within individual cells and enabling reconstruction of cell phylogenies^9–12^. However, these methods are difficult to scale and are hampered by high rates of genomic dropout and artifacts introduced during whole-genome amplification^9–12^. An alternative strategy is to detect somatic mutations directly from high-throughput single-cell assays such as single-nucleus RNA-seq and single-nucleus ATAC-seq, etc^13–16^, from transcribed regions and open chromatin regions of the genome. Besides variant calling, this approach can facilitate downstream functional annotation of the somatic mutations identified. Nonetheless, mutation detection in such data remains challenging. Recall and accuracy are limited by the lack of breadth and depth of coverage inherent to these technologies. Computational challenges exist to distinguish true variants from sequencing errors in sparse data and to delineate somatic mutations from germline polymorphisms^8,17,18^.

To address the variant calling challenges from SCS data, we and other groups have proposed several computational strategies. Among them, Monopogen^17^ leverages reference haplotype data and linkage disequilibrium (LD) to achieve accurate germline single nucleotide variant (SNV) detection and further uses detected neighboring germline SNVs as anchors to statistically delineate somatic SNVs, which have differential allelic segregation patterns across cells. SComatic^18^ distinguishes somatic from germline variants by exploiting the principle that germline variants should be present in all cell types, whereas somatic mutations are confined to restricted lineages with distinct gene expression patterns. While SComatic requires pre-partitioning of cells (e.g., based on RNA expression profiles) and performs somatic mutation calling at cluster level, Monopogen does not assume any pre-partitioning of the cells and can detect somatic mutations present in multiple RNA expression clusters. In this study, to evaluate the feasibility of somatic mutation calling from SCS data, we generated synthetic data by admixing HapMap cell lines from the SMaHT consortium to benchmark five variant calling approaches and highlight the potential advantages and challenges of performing somatic mutation detection using these technologies.

## Results

### Benchmarking sample and experiment design

To evaluate somatic variant detection from single-cell sequencing, a cell line mixture was produced using six HapMap samples at varying proportions, ranging from 0.5% up to 83.5% (**Figure 1A**). The sample that takes up the highest proportion in the cell-line mixture, HG005, serves as the germline background, whereas the other five samples serve as somatic clones. Single-nucleus RNA-seq (snRNA-seq) and single-nucleus ATAC-seq (snATAC-seq) libraries of this HapMap mixture were generated respectively and processed through the sequencing and the 10x Genomics Cell Ranger analysis pipeline. We obtained 12,170 estimated cells from snRNA-seq, with 36,585 mean reads per cell and 3,485 median genes per cell. For snATAC-seq, we got 10,963 estimated cells with 14,217 median high-quality fragments per cell. Matched PacBio long-read whole genome sequencing (WGS) data were also generated on each cell line, and variants detected from individual samples are compiled to serve as the ground truth. For this analysis, we focused on WGS data with snRNA-seq or snATAC-seq coverage with a minimum of 4 reads, ensuring that all ground-truth variants were evaluated under reliable coverage depth support (see **Methods**). After the quality control process, 10,479 and 7,767 cells were profiled from the HapMap mixture by snRNA-seq and snATAC-seq, respectively. We performed comprehensive assessments of somatic mutation detection capabilities in snRNA-seq and snATAC-seq datasets using Monopogen and SComatic. To accommodate the specific requirements of each variant calling program and to ensure accuracy, small modifications were made to each experimental workflow (see Methods). The SComatic pipeline requires cell type annotation as input. In our case, because all the cells in admixture are derived from B lymphocytes with indistinguishable RNA expression profiles, we used their donor source as the cell cluster labels instead, creating an optimistic execution condition for SComatic (see **Methods**). We performed demultiplexing for the HapMap mixture utilizing the WGS data from each donor. The demultiplexed proportions of different samples were consistent with the designed proportions (**Figure 1B**), indicating successful demultiplexing.

**Figure 1.**
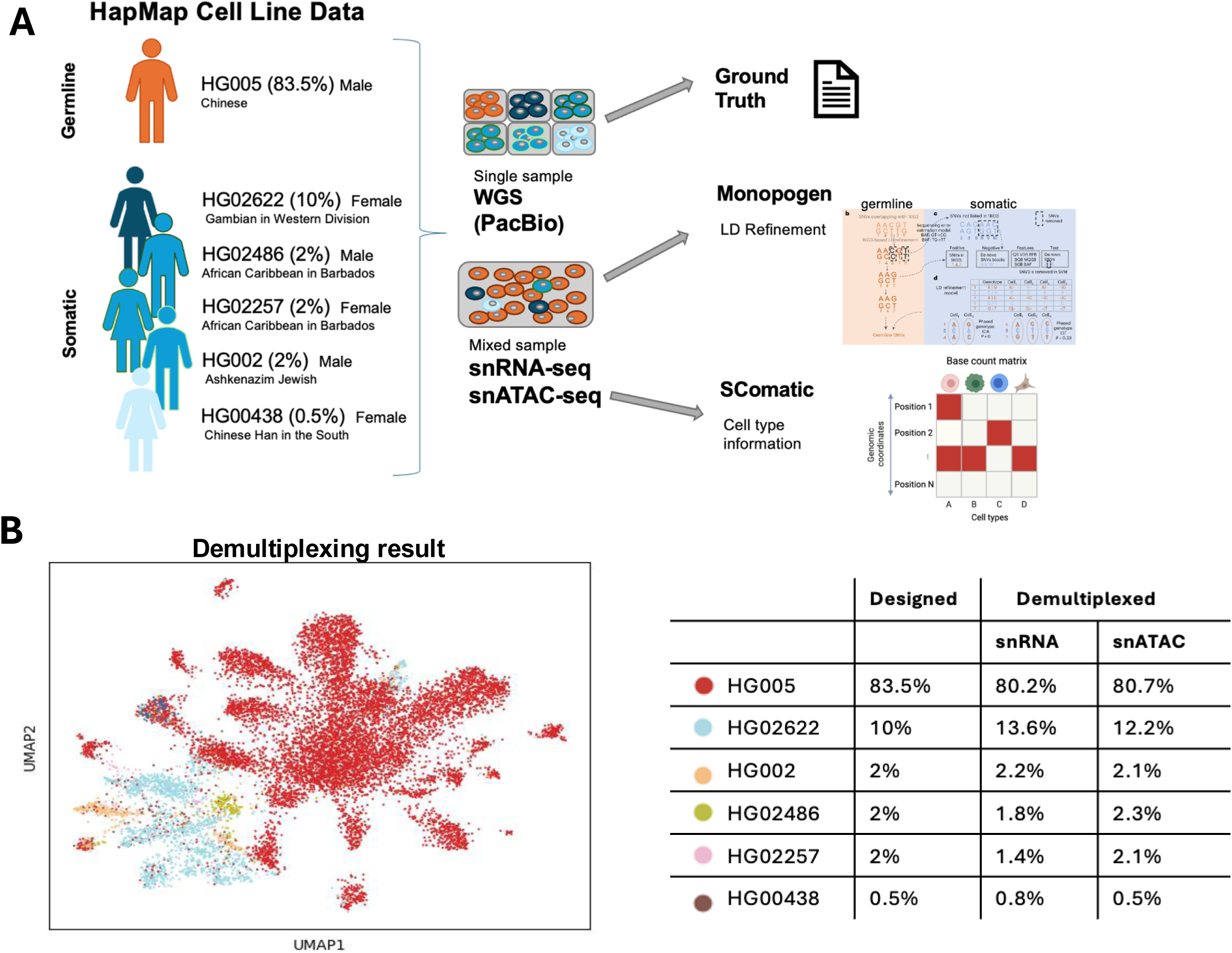

### Overview of variant calling from single-cell sequencing

For snRNA-seq data, Monopogen detected 119,945 SNVs in total, with 105,936 SNVs as germline and 14,009 SNVs as somatic (based on Monopogen’s classification rule). Out of the 119,945 SNVs, 111,906 (93.30%) of which are in the ground truth. By the design of the experiment, all of the germline variants called by Monopogen are in the ground truth. Out of the 14,009 somatic SNVs, 5,972 (42.63%) are in the ground truth somatic samples. In comparison, SComatic identified 31,655 SNVs, of which 26,369 SNVs were assigned to germline variants, and 5,286 SNVs were assigned to somatic mutations (based on SComatic’s classification rule). Out of the 31,655 SNVs, 23,537 (74.35%) of which are in the ground truth (**Table 1**, **Figure 3C**). Out of the 26,369 germline SNVs, 19,206 were in the ground truth (72.84%). Out of the 5,286 somatic variants, 4165 variants were in the ground truth (78.79%).

**Table 1.** Total SNVs detected by Benchmarking Methods.

| Total SNVs | snRNA-seq |  |  |  |  | snATAC-seq |  |  |  |  |
| --- | --- | --- | --- | --- | --- | --- | --- | --- | --- | --- |
|  | Monopogen | SComatic | DeepVariant | Cellsnp-lite | Mutect2 | Monopogen | SComatic | DeepVariant | Cellsnp-lite | Mutect2 |
| #SNVs | 119,945 | 31,655 | 209,942 | 118,668 | 0 | 2,165,773 | 105,217 | 2,692,151 | 1,063,453 | 78,567 |
| #In Ground Truth | 111,908 | 23,537 | 109,444 | 97,715 | 0 | 2,157,928 | 99,208 | 2,199,603 | 1,004,515 | 56,713 |
| #Not in Ground Truth | 8,037 | 8,118 | 100,498 | 20,953 | 0 | 7,845 | 6,009 | 492,548 | 58,938 | 21,854 |
| Precision | 93.30% | 74.35% | 52.13% | 82.34% | 0 | 99.64% | 94.29% | 81.70% | 94.46% | 72.18% |

Note: *SNV* Single Nucleotide Variant.

A significantly larger number of variants are detected in the snATAC-seq data compared with the snRNA-seq data. Monopogen detected 2,165,773 SNVs in total, with 2,125,033 germline SNVs and 40,740 somatic SNVs. Out of the 2,165,773 SNVs, 2,157,928 (99.64%) of which are in the ground truth. Out of the 40,740 somatic SNVs, 26,893 (66.01%) are in the ground truth somatic samples. In comparison, a much smaller number of variants were identified by SComatic, totaling 105,217 SNVs, of which 93,940 SNVs were assigned to the germline (HG005) sample, and 11,277 SNVs to somatic (non-HG005) samples. Out of the 105,217 SNVs, 99,208 (94.29%) of which are in the ground truth (**Table 1**, **Figure 3C**). Out of the 93,940 germline SNVs, 88,436 were in the ground truth (94.14%). Out of the 11,277 somatic variants, 10,493 were in the ground truth (93.05%).

Additional benchmarking results for DeepVariant, Cellsnp-lite and Mutect2 were presented in **Table 1 and Figure 3C**. Due to specific calling strategies, we used bulk WGS-identified variants as a prior for Cellsnp-lite calling (see **Methods**), and Mutect2 resulted with no mutations passing quality filtering for snRNA-seq (see **Discussions**).

### Characterization of read coverage and allelic frequency of variants

Across the genomic loci where variants are detected by the snRNA-seq and snATAC-seq, a more broadly distributed read depth pattern was found in snRNA-seq data, reflecting the wide dynamic range of RNA expression levels. In contrast, read depths of snATAC-seq data were more tightly clustered around a narrow range, indicating relatively similar coverage across open chromatin regions (**Supplementary Figure 1A-B**). Similar read depth patterns were observed from loci overlapped with the WGS ground truth variants (**Supplementary Figure 1C-D**), and from the somatic mutations identified by Monopogen, indicating the mutations detected by the computational strategy recapitulate the coverage characteristics of the true variants in both datasets (**Supplementary Figure 1E-F**).

We further calculated the detected variant allele frequency (VAF) from the single-cell sequencing data and compared it with the designed VAF. To avoid the impact of variants that are common in multiple samples on the VAF calculation, here we included only variants that are unique to a single sample. The detected VAFs of Monopogen somatic mutations were shown in **Figure 2**. Samples HG02622, HG02257, and HG00438 represented the designed mixed proportion of 10%, 2%, and 0.5%. Because variant calling was restricted to heterozygous SNVs in diploid regions, the expected VAFs should be 5%, 1%, and 0.25%, respectively. For both snRNA-seq and snATAC-seq data, the detected VAF peaks appeared around 0.05 for HG02622 (**Figure 2A-B**), 0.01 for HG02257 (**Figure 2C-D**), and 0.0025 for HG00438 (**Figure 2E-F**), which are highly consistent with the expectation. Similar VAF patterns were also found in the SComatic results (**Supplementary Figure 2**).

**Figure 2.**
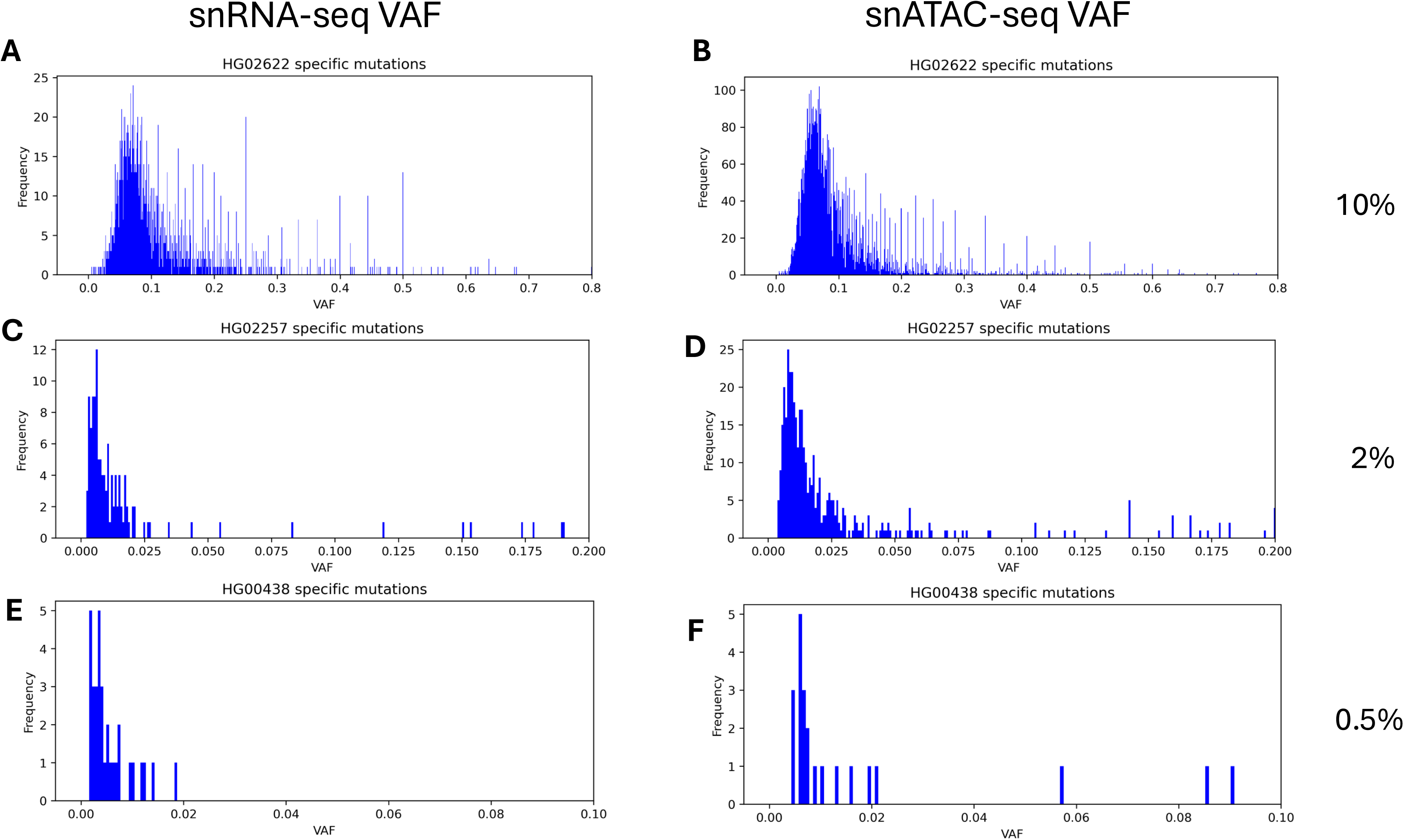

### Identification of donor-specific variants

To better assess the ability of the two tools in identifying somatic mutations from subclones, we evaluated their performance in recovering donor-specific variants. **Tables 2**–**3** showed the donor-specific comparison for snRNA-seq and snATAC-seq mutation between the two methods. In snRNA-seq data, Monopogen identified 4,523 SNVs in HG02622 sample, 2,213 SNVs in HG02486 sample, 2,059 SNVs in HG02257 sample, 1,505 SNVs in HG002 sample, and 1,054 SNVs in HG00438 sample. While Monopogen could not directly provide SNV donor-origin assignments, SComatic assigned SNVs to five samples with highly variable accuracy (with demultiplexing information provided). While the method showed reasonable precision for HG02486 (322/386, 83.4%) and HG02622 (3,173/3,875, 81.9%), it demonstrated notably lower precision for HG02257 (305/504, 60.5%) and especially HG00438 (82/175, 46.9%). More critically, SComatic exhibited low recall across all samples, detecting only a small fraction of the true variants present in the ground truth.

**Table 2.** Donor-specific variant calling results of snRNA-seq data.

| #SNV | %Mix | WGS | snRNA-seq |  |  |  |  |  |
| --- | --- | --- | --- | --- | --- | --- | --- | --- |
|  |  | Ground Truth | Monopogen<br>(germline) | Monopogen<br>(somatic) | SComatic | SComatic<br>(disregard<br>specific<br>samples) | DeepVariant | Cellsnp-lite |
| HG005 | 83.5 | 123,186 | 105,936 | 529 | 19,206 | 19,332 | 97,473 | 94,410 |
| HG02622 | 10 | 84,118 | 0 | 4,523 | 3,173 | 3,195 | 11,122 | 3,358 |
| HG02486 | 2 | 83,115 | 0 | 2,213 | 322 | 1,138 | 5,919 | 2,828 |
| HG02257 | 2 | 83,488 | 0 | 2,059 | 305 | 1,133 | 5,755 | 2,769 |
| HG002 | 2 | 52,348 | 0 | 1,505 | 280 | 711 | 3,875 | 2,122 |
| HG00438 | 0.5 | 41,535 | 0 | 1,054 | 82 | 475 | 2,903 | 1,659 |
| Not in Ground<br>Truth | - | - | 0 | 8,037 | 8,118 | 8,118 | 100,498 | 20,953 |

Note: *SNV* Single Nucleotide Variant.

**Table 3.** Donor-specific variant calling results of snATAC-seq data.

| #SNV | %Mix | WGS | snATAC-seq |  |  |  |  |  |  |
| --- | --- | --- | --- | --- | --- | --- | --- | --- | --- |
|  |  | Ground Truth | Monopogen (germline) | Monopogen (somatic) | SComatic | SComatic (disregard specific samples) | DeepVariant | Cellsnp-lite | Mutect2 |
| HG005 | 83.5 | 2,389,008 | 2,125,033 | 5,990 | 88,436 | 88,516 | 2,029,816 | 974,394 | 18,565 |
| HG02622 | 10 | 1,566,981 | 0 | 22,615 | 8,936 | 9,040 | 128,338 | 31,680 | 28,623 |
| HG02486 | 2 | 1,514,977 | 0 | 13,897 | 765 | 2,709 | 113,328 | 26,169 | 13,963 |
| HG02257 | 2 | 1,496,913 | 0 | 12,990 | 300 | 2,497 | 105,311 | 25,760 | 12,064 |
| HG002 | 2 | 963,527 | 0 | 9,832 | 473 | 1,493 | 80,198 | 19,997 | 7,231 |
| HG00438 | 0.5 | 770,117 | 0 | 6,212 | 16 | 1,151 | 51,305 | 14,668 | 3,937 |
| Not in Ground Truth | - | - | 0 | 7,845 | 6,009 | 6,009 | 492,548 | 0 | 21,854 |

Note: SNV Single Nucleotide Variant.

Almost 20 times more variants were identified by snATAC-seq than by snRNA-seq data, demonstrating the broader genome coverage of chromatin accessibility regions. In snATAC-seq data, Monopogen identified 22,615 SNVs from HG02622 sample, 13,897 SNVs from HG02486 sample, 12,990 SNVs from HG02257, 9,832 SNVs from HG002 sample, and 6,212 SNVs from HG00438 sample. Similar observations could be made for the variants called by SComatic as in the snRNA data. It assigned SNVs to five samples with consistently high precision. The method showed strong performance for HG02622 (8,936/9,640, 92.7% precision), HG02486 (765/801, 95.5%), HG002 (473/491, 96.3%), and HG02257 (300/320, 93.8%), indicating that most called variants were genuine. However, performance dropped notably for HG00438 (16/25, 64.0% precision). Despite the high precision, SComatic demonstrated low recall across samples, with substantially fewer variants called compared with the ground truth.

For variants from HG005 (defined as germline), Monopogen detected 86.00% (snRNA-seq) and 88.95% (snATAC-seq) respectively in the ground truth set, while SComatic detected 15.69% (snRNA-seq) and 2.42% (snATAC-seq). For other donor-specific somatic mutations in the ground truth set, Monopogen called about 2 times (snRNA-seq) and 5 times (snATAC-seq) more mutations than SComatic.

We performed a 3-way comparison of variants identified respectively by Monopogen, SComatic, and ground truth (**Figure 3A-B**). While the majority of ground truth variants were missed by both callers due to limited sequencing coverage, Monopogen demonstrated substantially higher concordance with ground truth compared with that of SComatic. The somatic mutation recall percentages from the WGS ground truth were presented in **Figure 3D-E**. The somatic mutation recall percentages from the WGS ground truth were presented in **Figure 3D-E**. Monopogen performed better than SComatic in both the snRNA-seq and the snATAC-seq data (average recall 3.18% vs 1.81% in the snRNA-seq data, 1.01% vs 0.25% in the snATAC-seq data). Interestingly, for both Monopogen and SComatic, the average recall percentages were consistently maintained for clones at both 2% and the 0.5% abundance (Monopogen: 2.66% vs 2.54% in the snRNA-seq data, 0.94% vs 0.81% in the snATAC-seq data; SComatic: 1.36% vs 1.14% in the snRNA-seq data, 0.17% vs 0.15% in the snATAC-seq data), indicating that somatic mutations in the least abundant (0.5% abundant) clone can be robustly detected, albeit at a low recall rate.

**Figure 3.**
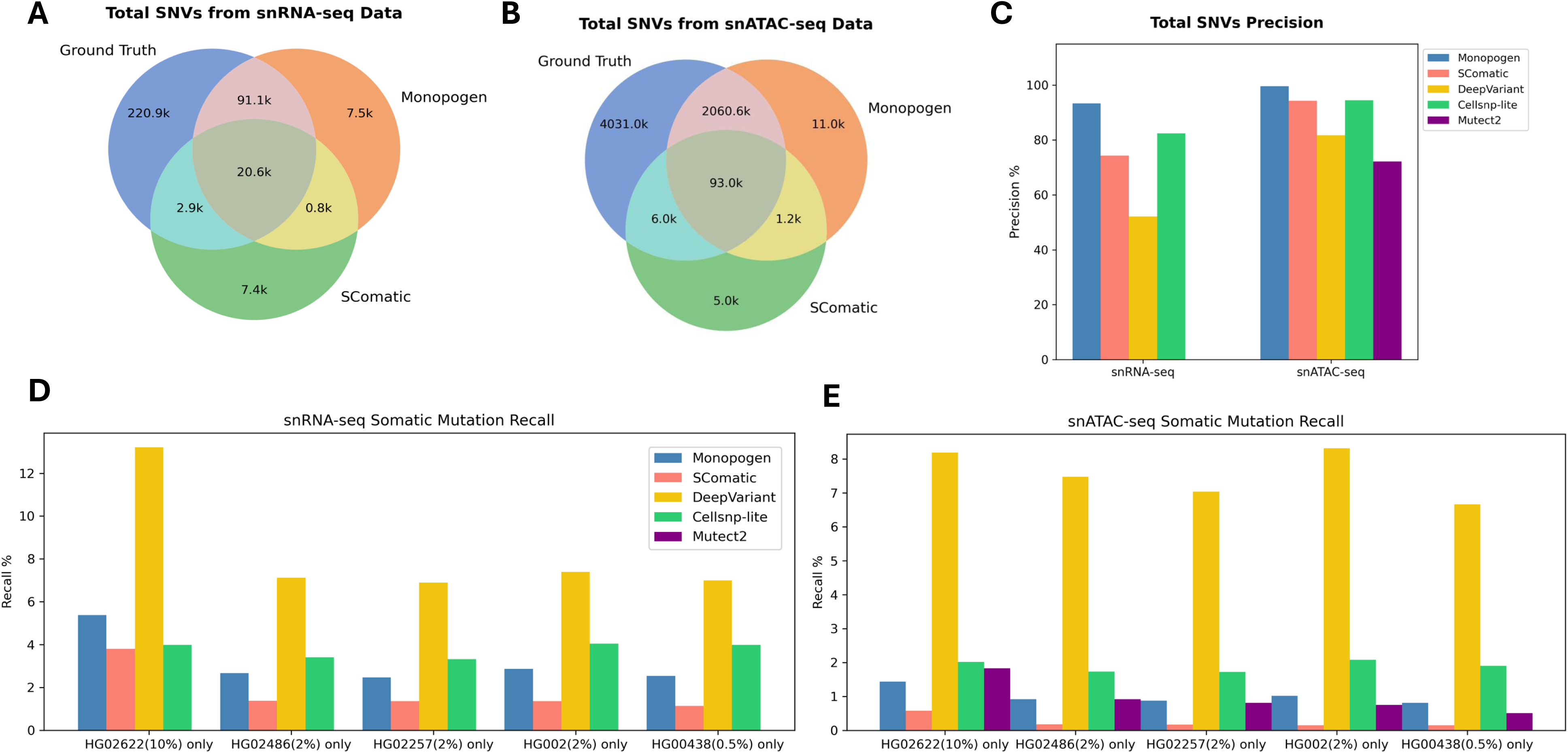

The comparison for other benchmarking methods was also presented in **Tables 2**–**3 and Figure 3D-E**. DeepVariant showed overall higher recall rates with lower precision compared with other methods. The recall rates were almost consistently maintained for different sample proportions. We have discussed the different calling strategies and their applications in the **Discussion** section.

### Distribution of donor-specific variants

Examination of the donor-specific mutation calls demonstrated substantial variation in how each method distributed mutations across the donor population. Both Monopogen germline module and SComatic identified more germline (HG005) SNVs than somatic (non-HG005) SNVs, while Monopogen somatic module identified more somatic mutations than SComatic (**Tables 2**–**3**). An important advantage of SCS technology is the ability to obtain the cellular origin of the detected variants. Given the design of the SComatic program, which explicitly assigns variants to donors, we performed a detailed analysis of its donor assignment accuracy by comparing mutations called without considering donor source against those assigned to specific donor sources. When comparing the two numbers, we found that donors with lower cellular abundance in the mixed population (non-HG005) were more likely to have their mutation misassigned to other donors (**Tables 2**–**3**). For example, in snRNA-seq data (**Table 2**), for donor HG02486, 1,138 true somatic variants were present in this donor, but SComatic assigned only 322 of them to this donor (28.3%), with shared variants assigned to other donors. Similar patterns were observed for other donors and in snATAC-seq data, which suggests that one of the major limitations of the method is its inability to assign variants to more than one cell type.

To check if the clonal origins of the variants detected by Monopogen were correctly assigned via cellular barcodes, we used the demultiplexing results as a proxy ground truth to evaluate the cell-origin assignments. We included only donor-specific variants in the calculation to avoid multiple cell-origin assignments and applied more strict filtering (see **Discussion**). Significantly higher accuracy of variant source assignments was observed in the Monopogen results (**Figure 4**). The snATAC-seq data showed higher overall accuracy compared to the snRNA-seq data (94.5% vs 90.2%). The sample with a higher mixed proportion (HG02622, 10%) showed higher accuracy over the other 2% mixed samples (HG002, HG02486, and HG02257). HG00438 came with the lowest cell proportion (0.5%) and exhibited the largest variation in assignment accuracy. The major misassignment came from the HG005 sample (with around 10% misassigned cell origins). Overall, the single-cell sequencing and mutation calling technology demonstrated more than 80% of mutation cell-origin assignment accuracy for most donors in both RNA and ATAC modalities.

**Figure 4.**
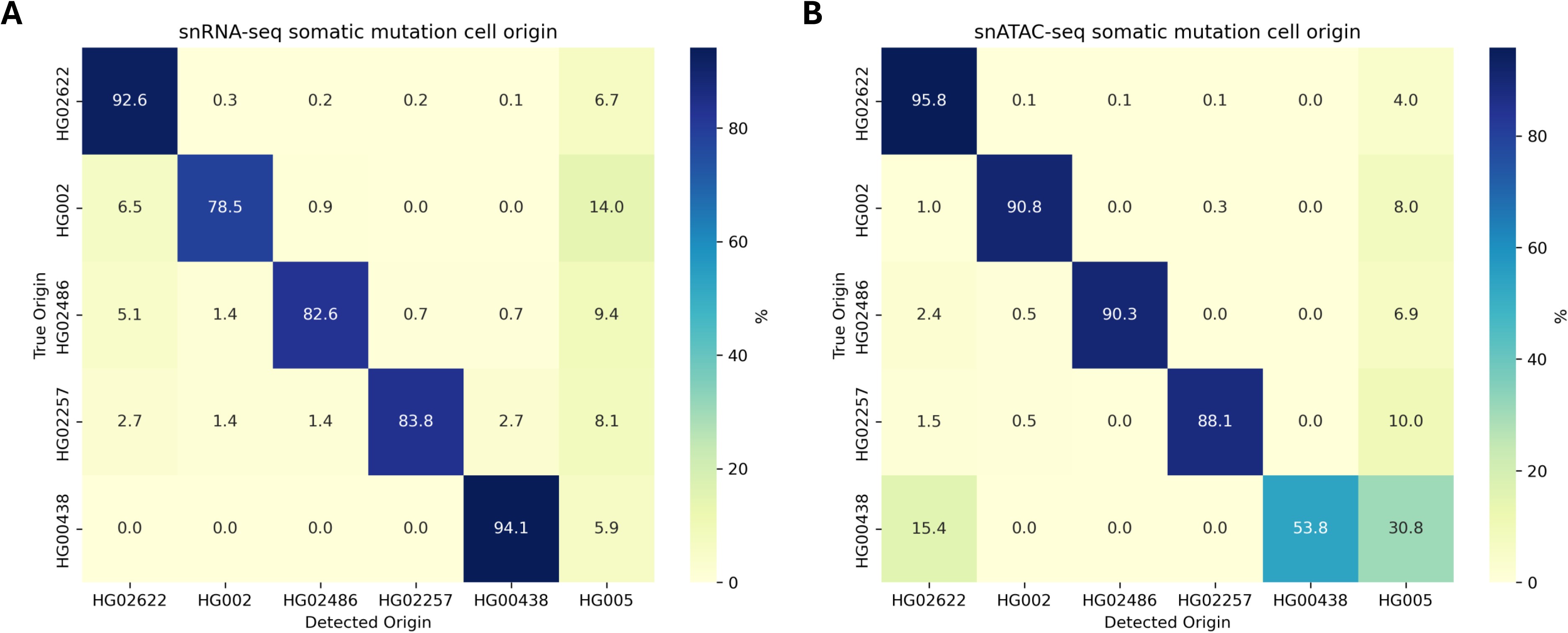

### Effect of Total Variant Read Depth on Variant Calling

To simulate real-world sequencing variability, we downsampled the full dataset to a range of sequencing depths and evaluated precision and recall across variants with varying coverage levels (see **Methods**). For total variants called, both modalities exhibited a modest decrease in precision and a corresponding increase in recall as sequencing depth increased (**Figure 5**). This trade-off likely reflects the detection of additional low-confidence variants at higher depths, which inflates recall at the cost of precision.

**Figure 5.**
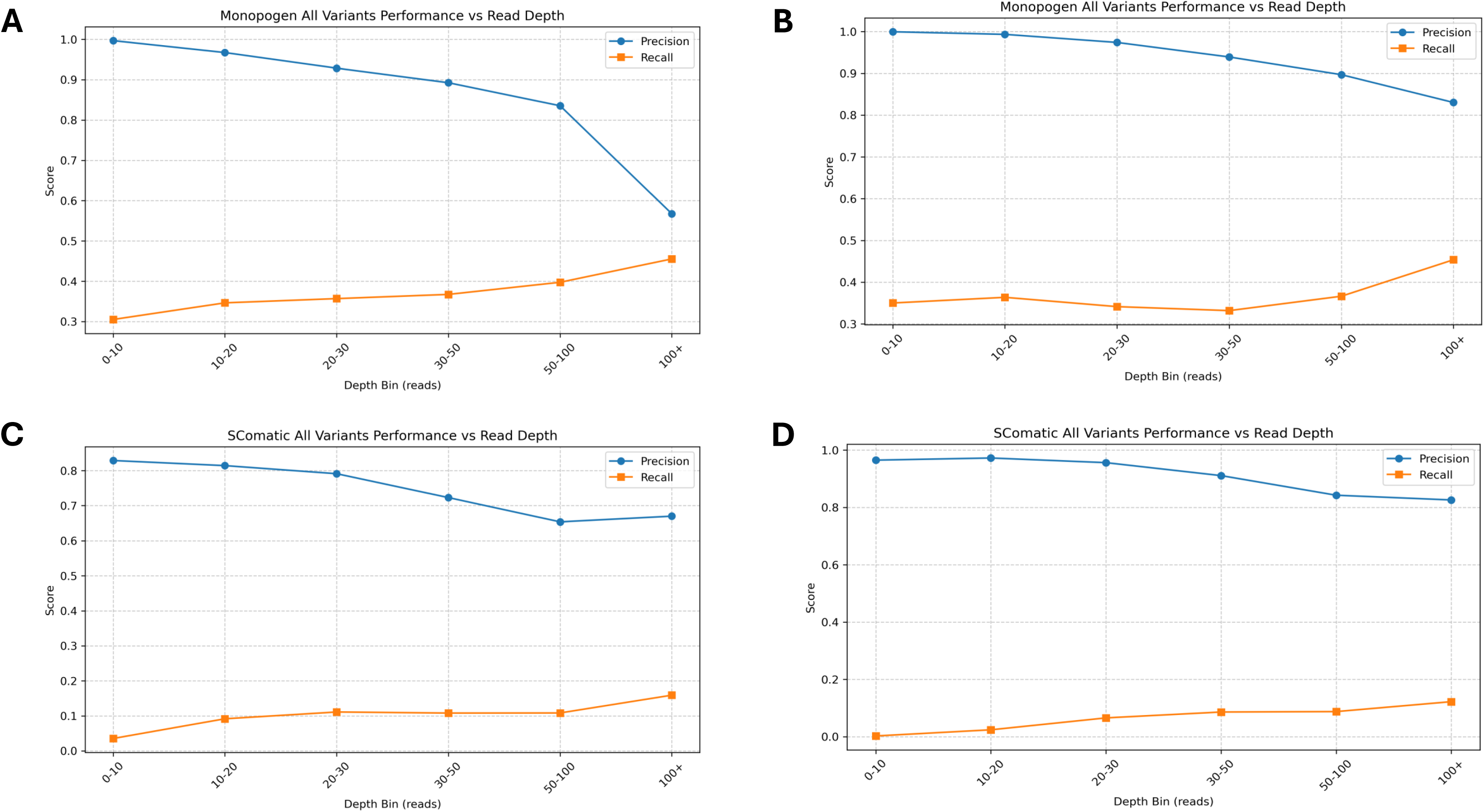

When stratifying by variant class, germline and somatic variants displayed distinct depth-dependent behaviors. In both modalities, germline variant recall peaked at an intermediate sequencing depth, suggesting that beyond a certain threshold, additional reads do not improve, and may marginally reduce, the recovery of true germline variants. In contrast, somatic variant calling benefited more consistently from increased depth, with both precision and recall improving monotonically, consistent with the expectation that low-allele-frequency somatic variants require greater coverage for reliable detection (**Supplementary Figures 3-4**).

In conclusion, our experiments indicate that it is feasible to sensitively detect low abundance (0.25% allele fraction) somatic mutations from snRNA-seq and snATAC-seq data, despite the low recall rate resulting from restricted sequencing coverage. Monopogen has better precision and recall for somatic mutation detection compared with SComatic. The somatic mutation detection rate remains quite stable at lower percentage samples. Thus, SCS could be used as a complementary approach to other methodologies for somatic variant detection in the normal tissues.

## Methods

### Sample Collection

B-Lymphocyte lymphoblastoid cell line samples from six donors were mixed at predetermined ratios and subjected to snRNA and snATAC sequencing. WGS data were also generated for each individual donor. In the HapMap mixture, sample HG005 represented the highest proportion (83.5%), and its variants were treated as germline mutations in downstream analyses. The variants from the rest of the donors were treated as somatic mutations in downstream analyses. Additional details about this HapMap mixture are in the benchmarking paper of the SMaHT Network^19^.

### Single Cell Sequencing Data Collection

The library preparation and sequencing of single-nuclei cDNA were carried out following the manufacturer’s protocols (https://www.10xgenomics.com). To obtain single-cell GEMS (Gel Beads-In-Emulsions) for the reaction, the single-nuclei suspension was loaded onto Chromium X. The pair-end library for single nuclei RNA-seq was prepared with the Chromium GEM-X Single Cell 5’ Reagent Kits v3 (10x Genomics), while the pair-end library of single nuclei ATAC-seq was prepared with the Chromium Next GEM Single Cell ATAC Reagent Kits v2 (10x Genomics). The constructed libraries were subsequently sequenced on an Illumina Novaseq 6000 (https://www.illumina.com).

### Data Preprocessing

Raw sequencing reads from snRNA-seq and snATAC-seq were processed using the 10x Genomics Cell Ranger pipeline (cellranger-7.2.0 for RNA-seq, cellranger-atac-2.1.0 for ATAC-seq, with GRCh38-2020-A reference genome). Quality control was conducted using an in-house developed computational pipeline. For snRNA-seq, we started from the raw count feature matrices of Cell Ranger and filtered cells with UMI counts below 500, gene counts below 300, and mitochondria fraction above 5%. We then used SoupX to purify the transcriptome measurement by subtracting the background transcripts. DoubletFinder further detected the potential doublets and retained the clean count feature matrices for singlets. For snATAC-seq, we used ArchR (v1.0.3) to filter doublets.

### Whole Genome Sequencing Data Analysis

To establish a genomic baseline, WGS was conducted on the 6 samples, providing comprehensive variant profiles for reference and comparative analyses. WGS libraries were prepared using the PacBio SMRTbell Prep Kit 3.0 and sequenced on the PacBio Revio sequencer to an average coverage of 100×. Sequencing reads were aligned to the human reference genome GRCh38 using pbmm2 (v.13.1). Variants were called using Clair3 (default parameters).

The ground truth for single-cell comparison was established by treating all variants from HG005 as germline variants. Somatic variants were defined as those present in each of the other samples’ WGS data that did not overlap with HG005. For the purpose of comparison, we retained only variants located in RNA/ATAC-covered regions with a minimum of four supporting RNA/ATAC reads.

### Single Cell Variant Calling from Monopogen

#### Preprocessing alignment file

Variant calling was performed using Monopogen with input VCFs generated from Samtools to pileup the reads and BCFtools for preliminary filtering. During preprocessing, reads with mapping quality below 20 and bases with quality scores below 20 were excluded from analysis. Read depth was recorded for each position, with the maximum depth threshold set to 10,000,000 to prevent down-sampling. Sites containing non-canonical reference alleles or symbolic entries were filtered out. Multiallelic variants were decomposed into biallelic representations and left-normalized against the reference genome to ensure consistent variant representation.

#### Germline mutation calling

The resulting high-quality, normalized VCFs served as input for Monopogen’s downstream processing pipeline. Monopogen employed a template-matching strategy to distinguish reference and alternative alleles on a per-cell basis, utilizing cell barcodes and UMI information to mitigate amplification artifacts inherent in single-cell sequencing data. For this experiment, a germline reference panel was established using variants from the 1000 Genomes Project that overlapped with the HG005 WGS data. This reference panel was then used to phase the raw alignments and derive the germline genotypes for the single-cell data.

#### Somatic mutation calling

To distinguish somatic from germline variants, Monopogen integrated multiple complementary features beyond raw allele counts. Candidate variant sites were initially evaluated against expected allele frequency distributions under germline heterozygosity assumptions. Sites exhibiting deviations consistent with non-germline patterns were flagged as putative somatic events. These preliminary somatic calls underwent further refinement using linkage disequilibrium (LD) structure analysis. True germline variants are expected to co-segregate with nearby alleles according to established LD patterns, whereas somatic variants typically disrupt these linkage relationships.

### Single Cell Variant Calling from SComatic

The SComatic pipeline requires cell type information for each cell as input data. In this experimental design, although these cells constitute a homogenous population of B-lymphocyte origin, we utilized donor specific variants to simulate cell type specific variants. The demultiplexing results, which identify the donor source of each cell, were used as a proxy for cell type labels. While different cell types from a single donor share nearly identical genomic sequences, using multiple donors introduces much more unique germline variants. This donor-derived genomic diversity provides a highly powered experimental condition for SComatic.

Demultiplexing was conducted using popscle demuxlet, with WGS variants as the reference file. Downstream analysis followed the SComatic pipeline steps 1 to 4.1, using the default beta binomial parameters and setting parameters “--min_dp”, “--min_cc”, “--min_cov”, “--min_cells”, “--min_ac_cells”, “--min_ac_reads” all to one to include as many variant candidates as possible. We further filtered the candidates with a minimum of four supporting RNA/ATAC reads.

### Single Cell Variant Calling from DeepVariant

Variant calling was performed using DeepVariant v1.10.0^20–21^. For snRNA-seq data, the RNAseq mode was applied to account for splicing-induced alignment artifacts. For snATAC-seq data, WES mode was used given the open chromatin capture strategy. In both cases, the small model was disabled, and variant calling was performed across entire chromosomes rather than restricted intervals. Raw VCF files were subsequently processed by splitting multiallelic sites into biallelic records, retaining only SNV sites, and restricting to variants with a PASS filter designation.

### Single Cell Variant Calling from Cellsnp-lite

We performed variant calling using Cellsnp-lite v1.2.3 (htslib v1.22.1)^22^. The input BAM file was derived from Cell Ranger output, and candidate variants were defined by a WGS-derived VCF (PacBio for the HapMap mixture sample) filtered to sites with a minimum read depth of 2. We set the minimum read count to 4 and applied no minimum minor allele frequency threshold.

### Single Cell Variant Calling from Mutect2

Somatic variant calling was performed using Mutect2 in tumor-only mode (GATK v4.6.2.0)^23–26^. Prior to variant calling, BAM files were preprocessed through a sequential pipeline: duplicate reads were flagged with MarkDuplicates, read groups were assigned with AddOrReplaceReadGroups, and reads were reordered to match the reference dictionary using ReorderSam. For RNA-seq inputs, reads spanning splice junctions were split using SplitNCigarReads. Base quality scores were then recalibrated with BaseQualityScoreRecalibrator (BQSR). Somatic variants were called with Mutect2 and subsequently filtered using FilterMutectCalls with default parameters.

### Comparison of Mutations Detected in snRNA-seq, snATAC-seq and WGS data

For performance comparison, mutations were grouped into three categories: (1) true positives (variants detected in both the single-cell dataset and the WGS data); (2) false positives (variants identified in the single-cell dataset but absent from the WGS data); and (3) false negatives (variants present in the WGS data within regions covered by the single-cell assay but undetected by the single-cell method). Precision for each method was then calculated as precision = True Positives / (True Positives + False Positives). Recall for each method was calculated as recall = True Positives / (True Positives + False Negatives).

Given the different assumptions between the two methods, we further compared their performance at the individual donor level. For variants called by Monopogen, we examined the number of variants identified in the germline and somatic calling steps, respectively. For variants called by SComatic, we compared the donor-specific variants identified based on the demultiplexing-derived donor labels with those detected without considering the demultiplexed labels.

### Effect of Total Variant Read Depth on Precision and Recall

Variants were stratified by sequencing coverage and binned to assess precision and recall across depth ranges. Coverage at each variant position was obtained using bedtools coverage applied to the input BAM file. Variants were assigned to the following depth bins: [0–10), [10–20), [20–30), [30–50), [50–100), and ≥100 reads, chosen to reflect the range of coverage commonly observed in single-cell sequencing data. Precision and recall were calculated independently within each bin.

For germline and somatic specific analyses, variants were categorized as follows. Germline variants were defined as those called by the Monopogen germline module or assigned to sample HG005 by SComatic, with ground truth derived from the WGS callset of HG005. Somatic variants comprised those called by the Monopogen somatic module or assigned to the remaining samples in SComatic, with ground truth defined as WGS variants absent in HG005.

## Discussion

Here we present a benchmark analysis evaluating the feasibility of detecting somatic mutations from single-cell RNA-seq and ATAC-seq data. While numerous benchmarking studies have evaluated somatic mutation detection using bulk sequencing data^27–30^, assessments at the single-cell level remain limited. Existing evaluations of single-cell somatic variant calling have primarily focused on full-length SMART-seq transcriptomic data^31^ or single-cell DNA sequencing datasets^32^. More recently, Matthew et al. systematically assessed somatic mutation calling using scRNA-seq and snATAC-seq data^33^; however, their work relied predominantly on bulk-oriented somatic variant callers, and comprehensive evaluation of single-cell–specific somatic calling methods remains largely absent. Our benchmarking evaluation employs a more robust experimental design using an ideal cell mixture with predefined ratios and corresponding WGS from each donor. This design allows for a more rigorous assessment of whether the widely used snRNA-seq and snATAC-seq approaches can reliably identify mutations compared to bulk sequencing methods. With designed germline and somatic variants in the cell line experiment, our framework provides a comprehensive understanding of detection accuracy and recall for somatic mutation calling using the most advanced individual-cell level variant callers currently available^17,34^. In this study, we focused specifically on somatic SNV detection. Other forms of somatic variation, including indels, CNVs, and retrotransposition events, are also important but are beyond the scope of the current work. Detection of large-scale somatic genomic alterations from short-read single-cell sequencing data remains technically challenging due to the low sequencing coverage. Our analysis demonstrates that somatic variant detection from single-nucleus sequencing data is feasible even at low frequencies.

From the benchmark cell line admixture, we successfully identified somatic variants with allele frequencies as low as 0.5%, indicating that single-cell technologies can indeed capture rare mutational events within heterogeneous cell populations. The overall accuracy of variant calls was high, with precision rates exceeding 90% for most datasets, confirming that the detected variants represent genuine mutations rather than technical artifacts. When comparing the two modalities, snATAC-seq outperformed snRNA-seq in both genomic coverage breadth and detection accuracy, likely due to the more uniformed genomic distribution and high per-locus coverage compared with transcript-dependent coverage in snRNA-seq. The recall rates were consistently maintained for rare variants with expected allele frequencies lower than 2%, specifically in variant callers designed for handling single-cell data, ranging from 0.025 to 0.029 (Monopogen), 0.011 to 0.014 (SComatic) for snRNA seq, and 0.008 to 0.010 (Monopogen), 0.0015 to 0.0018 (SComatic) for snATAC-seq data. Although the WGS-independent methods show lower recall relative to the whole genome, the variants detected are predominantly from protein coding and gene regulatory regions, which are highly valuable for downstream functional genomic analysis. These findings establish single-nucleus sequencing as a viable platform for somatic variant detection while highlighting that adequate sequencing depth remains the critical bottleneck for achieving comprehensive variant discovery.

Having the synthetic cell-line mixture data is important in enabling objective assessment of the power of the SCS platforms and the variant calling algorithms. That said, our current experiment only allowed us to objectively examine variants presenting at greater than 0.25% allelic frequency. Given the informative results of our experiments, it is possible to further titrate the cell-lines to examine the potential of calling even lower frequency variants using SCS.

The primary limitation of single-nucleus sequencing for variant detection is limited recall due to insufficient sequencing coverage under the current standard 10x SCS data generation protocols. Unlike bulk methods that cover reads at each locus, single-cell approaches distribute reads across each cell, resulting in sparse per-locus coverage. We only achieved less than 1% recall for each donor despite analyzing over 10,000 cells. During benchmarking, we also found that site with adequate coverage (≥ 4 reads) achieved higher recall compared with the under-coverage site. The trade-off between cellular resolution and genomic coverage is the major constraint of the single-cell methods, indicating that increased sequencing depth could facilitate comprehensive mutation detection.

Another notable limitation of the current benchmarking framework is the use of a single cell type for performance evaluation. Variant calling performance in single-cell data is inherently influenced by cell-type-specific properties, including transcriptional activity, chromatin accessibility, and baseline mutation burden. For snRNA-seq-based callers, cells with higher transcriptional output provide greater per-site coverage across expressed loci, which may favorably bias recall estimates relative to transcriptionally quiescent cell types. Similarly, for snATAC-seq-based callers, the breadth and distribution of accessible chromatin regions vary substantially across cell types, affecting the genomic sites available for variant detection. Consequently, precision and recall estimates derived from a single cell type may not generalize across the full spectrum of cell types encountered in heterogeneous tissues or tumor microenvironments. Future benchmarking efforts should incorporate diverse cell types, including rare populations and cell states with distinct epigenomic and transcriptomic profiles, to provide a more comprehensive assessment of caller performance. Additionally, benchmarking across cell types with known differences in somatic mutation burden, such as neurons versus rapidly dividing progenitor cells, would offer valuable insight into how variant callers perform under biologically distinct conditions.

We found that variant calling is sensitive to the algorithm, with each algorithm exhibiting distinct strengths and weaknesses. For the SComatic method, we were able to recover somatic mutations with higher precision than Monopogen, while maintaining comparable recall for detecting low-frequency variants. However, SComatic showed critical limitations in variant assignment. The algorithm systematically assigned shared somatic mutations to a single donor, even when variants were present in different donors. For example, in **Table 2** and **Table 3**, the number of variants detected by disregarding specific samples is usually higher than when considering the sample assigned in SComatic. This one-to-one assignment could be problematic when cell type annotations were very stratified.

In contrast, Monopogen demonstrated robust sensitivity across both data modalities and identified somatic mutations with allele frequencies as low as 0.5% from snRNA-seq and snATAC-seq datasets (**Figure 2**, **Supplemental Figure 1**). A key strength of Monopogen is its ability to identify donor-specific mutations with high precision without any additional information. However, due to the germline filtering design of the program, which relies on the 1KG3 reference panel, Monopogen cannot distinguish germline mutations that are not in the reference panel but exhibit allelic imbalance from true somatic mutations. For example, in snRNA-seq data, 529 variants were identified as somatic variants in the pipeline, but they should be germline instead. With reference WGS from the donor, Monopogen could better distinguish non-1KG3 germline variants from true somatic mutations and would have improved performance in the somatic calling step by more effectively distinguishing true somatic mutations from sequencing artifacts. The SMaHT Network is generating deep WGS for all donors (100× long-read and 330× short-read coverage) and will also produce several donor-specific assemblies, providing opportunities to further improve the detection algorithm. It is worth noting that despite the clearly defined proportions of cell mixtures from different donors, our experimental design still has some limitations. The decision to use variants in HG005 as the germline variants overestimates the true number of germline variants present in the dataset, as HG005 also contains somatic mutations that are not present in the other five samples. This design could also affect the variant calling performance of Monopogen in two ways. First, we completely disregard variants in other samples that are present in the 1KG3 reference panel. Second, because Monopogen relies on population-level LD patterns, the cell mixture from different races and ethnicities might reduce the power of this method^35–36^. Our evaluation for SComatic also represents the most ideal scenario, as we used the demultiplexing results (based on the WGS reference) as a proxy for cell type input. Through our testing, we found that the accuracy of the resulting mutation calls varies highly depending on the cell type information provided.

DeepVariant is not designed for single-cell dataset. In this study, DeepVariant demonstrated more true positives detected with more false positives. In real datasets, it would be difficult to separate them. Cellsnp-lite utilizes WGS-identified variants as a prior to guide variant-to-cell assignment, which introduces a fundamental limitation: the method is constrained to variants already present in the WGS input and cannot detect de novo mutations arising independently in individual cells. In real datasets, low-allele-frequency variants may fall below the detection threshold of bulk WGS and therefore be absent from the input variant list entirely. This is particularly consequential for somatic evolution studies, where rare de novo mutations, undetectable by bulk sequencing, may represent early clonal events or driver alterations with direct biological relevance. On the other hand, this could be a good validation method to check candidate variants called from other sequencing methods on single-cell data.

Many somatic variant callers require matched tumor-normal pairs for reliable mutation detection. Among these, we selected Mutect2 as a benchmark representative, applying the GATK-recommended panel of normals to suppress germline and artifactual variants. However, because our analysis was performed on normal tissue in tumor-only mode, without a matched normal sample for direct comparison, variant detection was substantially compromised. In snRNA-seq data, no mutations passed quality filtering, likely due to the confounding influence of RNA editing events, which can produce systematic nucleotide changes that are indistinguishable from genuine somatic variants under this calling framework. In snATAC-seq data, Mutect2 detected substantially fewer variants than other benchmarked methods, with notably lower precision, consistent with the elevated false positive burden expected when running a tumor-normal caller in the absence of a matched normal.

We found that somatic variants can be assigned to their correct cellular source with high accuracy using single-nucleus sequencing data. Monopogen achieved more than 80% of assignment accuracy for most donors (**Figure 4**). For the variants that were misassigned, the error source could be: (1) the background contamination of ambient RNAs, which could contain variants from other cells and samples; (2) sequencing related false calls, including low base-call quality or systematic errors, RT/PCR-induced errors, mapping or alignment issues; (3) the demultiplexing errors, that the cell-origin ground truth could be wrong in this experiment design^33,35,36^. The major misassignment originated from the HG005 sample, possibly due to its highest cell proportion (83.5%). HG00438 had the lowest cell proportion (0.5%), and only a few somatic mutations were detected from this sample, which may have limited the robustness of its representation in the results. Here we filtered out called somatic mutations with less than two reads per cell or less than two cells detected with that mutation. It is likely that further fine-tuning of the filter criteria can lead to more reliable mutation calls. Overall, the high mutation cell-origin assignment accuracy indicated that with real tissue sample data, we could also trace back the specific cell type that contains that specific somatic mutation.

In conclusion, single-nucleus RNA-seq and ATAC-seq could enable somatic variant detection at single-cell resolution, with the ability to identify mutations as rare as 0.5% allele frequencies and accurately assign them to specific cell types. snATAC-seq demonstrated better performance than snRNA-seq, with broader genomic coverage and higher accuracy. However, limited recall due to sparse sequencing coverage is still a fundamental constraint. Future algorithmic improvements can be made on germline-filtering methods independent of reference panels and more accurate multi-donor variant assignments. Further improvement can be achieved by integrating with complementary approaches such as bulk whole-genome sequencing or targeted deep sequencing, leveraging its unique strength of cellular resolution while compensating for limited genomic coverage. This integrative strategy would enable robust characterization of clonal evolution and cell-type-specific mutational landscapes in heterogeneous tissues.

## Acknowledgements

This research is supported by the NIH Common Fund, through the Office of Strategic Coordination/Office of the NIH Director under awards UM1 DA058229. This work is made possible by DI3-0000000288 from the Chan Zuckerberg Initiative DAF, an advised fund of the Chan Zuckerberg Initiative Foundation.

## Author Contributions

R.C., K.C., and R.A.G. designed, planned, and supervised the project. H.D, S.V.B., and D.K. generated the sequencing data. C.M.G. and H.D. managed sequencing data transfer. R.L. and Z.W. analyzed the data and drafted the manuscript. J.D., R.C., and K.C. provided successive revisions of early drafts. All authors provided critical reviews of the manuscript.

## Data Availability

The datasets described in this study will be made available through dbGaP under the study accession numbers phs004193. More information about the SMaHT Network is available online at https://smaht.org/, about the SMaHT Data Portal at https://data.smaht.org/, and types of data generated by the Network at https://data.smaht.org/about/consortium/data.

