## Supplementary Figures for "Somatic variant detection in normal tissues from single-cell sequencing data"

### Slide 1
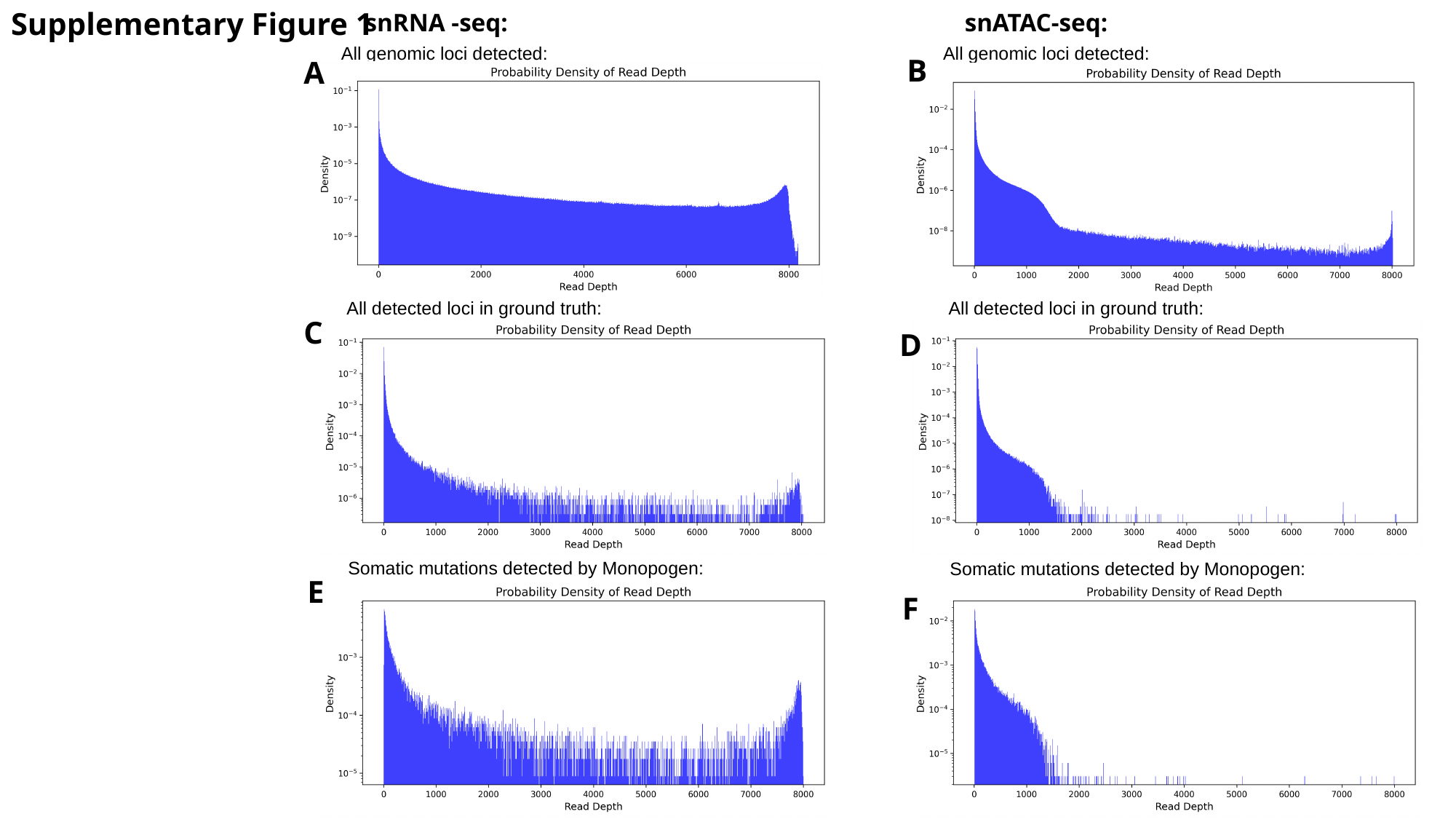

Supplementary Figure 1
snRNA -seq:
snATAC-seq:
All genomic loci detected:
All genomic loci detected:
B
A
All detected loci in ground truth:
All detected loci in ground truth:
C
D
Somatic mutations detected by Monopogen:
Somatic mutations detected by Monopogen:
E
F

### Slide 2
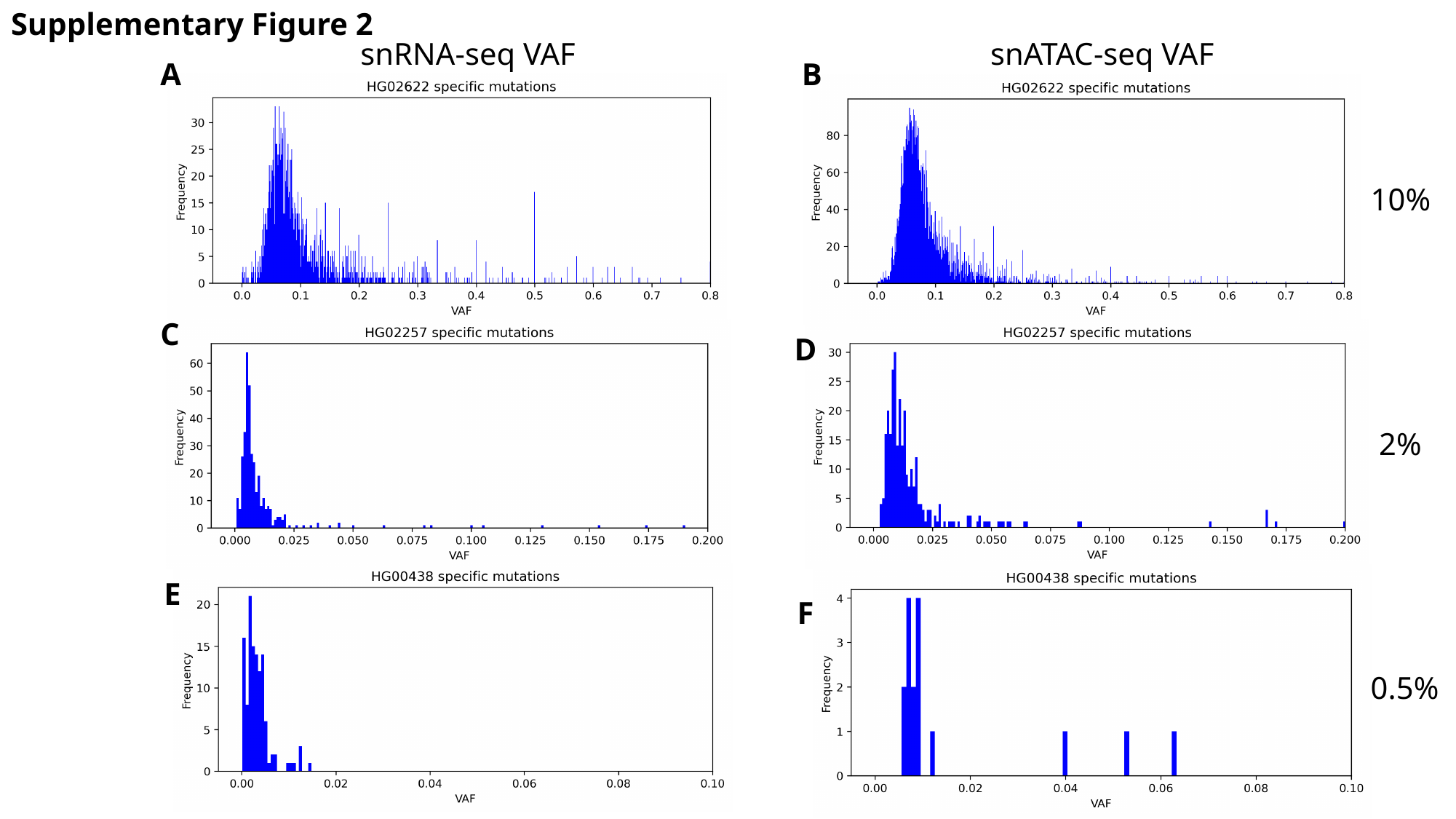

Supplementary Figure 2
snRNA-seq VAF
snATAC-seq VAF
A
B
10%
C
D
2%
E
F
0.5%

### Slide 3
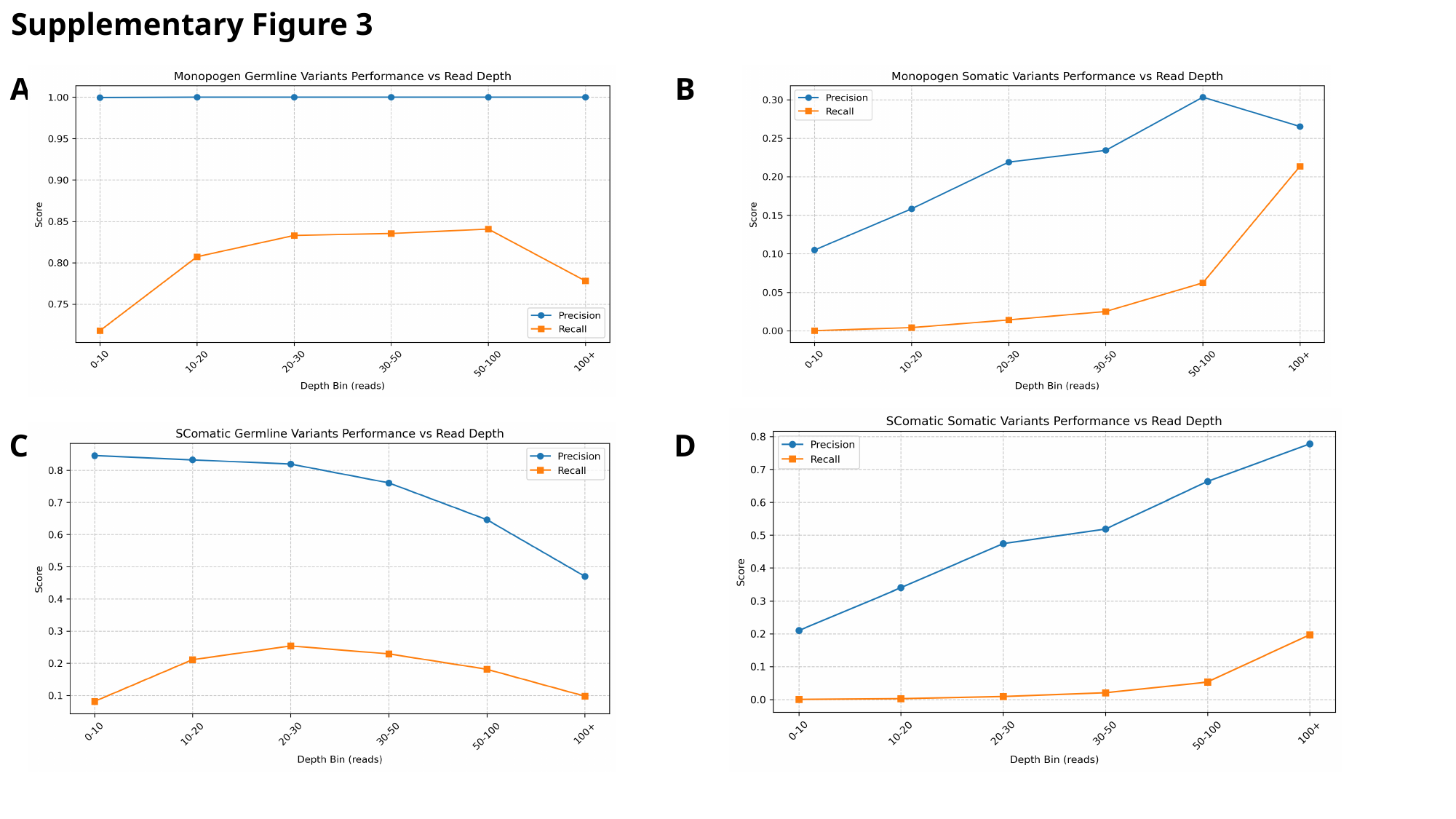

Supplementary Figure 3
A
B
C
D

### Slide 4
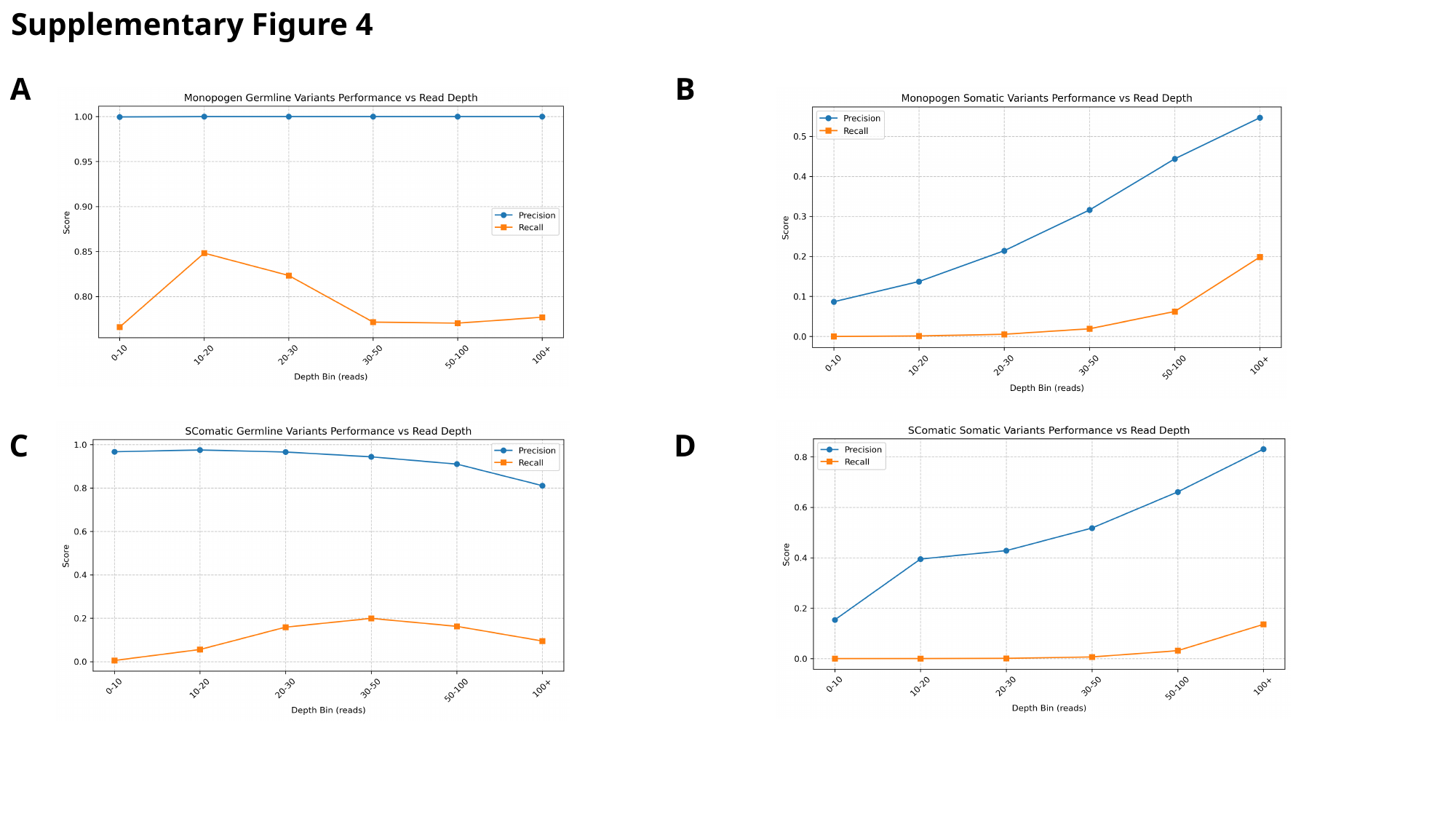

Supplementary Figure 4
A
B
C
D
